# *ITGA2* is a target of miR-206 promoting cancer stemness and lung metastasis through enhanced *ACLY* and *CCND1* expression in triple negative breast cancer

**DOI:** 10.1101/583062

**Authors:** Valery Adorno-Cruz, Andrew D. Hoffmann, Xia Liu, Brian Wray, Ruth A. Keri, Huiping Liu

**Affiliations:** Department of Pharmacology, Case Western Reserve University, Cleveland, OH, USA.; Department of Pharmacology, Northwestern University Feinberg School of Medicine, Chicago, IL, USA.; Cleveland Clinic Foundation, Cleveland, OH, USA.; Bioinformatic Core, Northwestern University Feinberg School of Medicine, Chicago, IL, USA.; Department of Genetics and Genome Sciences, the Division of General Medical Sciences-Oncology, Case Western Reserve University, Cleveland, OH, USA.; Department of Medicine, the Division of Hematology and Oncology, Northwestern University Feinberg School of Medicine, Chicago, IL, USA.; Lurie Comprehensive Cancer Center, Northwestern University Feinberg School of Medicine, Chicago, IL, USA.; Department of Pathology and Case Comprehensive Cancer Center, Case Western Reserve University, Cleveland, OH, USA.

## Abstract

Accumulating evidence demonstrates that cancer stemness is essential for both tumor development and progression, regulated by multi-layer factors at genetic, epigenetic and micro-environmental levels. However, how to target stemness-driven plasticity and eliminate metastasis remains one of the biggest challenges in the clinic. We aim to identify novel molecular mechanisms underlying stemness of triple negative breast cancer (TNBC) which frequently metastasizes to the visceral organs but lacks targeted therapies. Following our previous discovery of miR-206 as an epigenetic suppressor of tumorigenesis and metastasis, we now report that the integrin receptor CD49b-encoding *ITGA2* is an oncogenic target of miR-206 in TNBC. *ITGA2* knockdown abolished cancer stemness (mammosphere formation, pluripotency marker expression, and FAK phosphorylation), inhibited cell cycling, compromised migration and invasion, and thereby decreasing lung metastasis of TNBC. RNA sequencing analyses of breast cancer cells revealed that *ITGA2* knockdown inhibits gene expression essential for both classical integrin-regulated pathways (cell cycle, wounding response, protein kinase, etc) and newly identified pathways such as lipid metabolism. Notably, *ACLY*-encoded ATP citrate lyase is one of the top targets in CD49b-regulated lipid metabolism and *CCND1*-encoded Cyclin D1 represents regulation of cell cycle and many other pathways. ACLY, known to catalyze the formation of cytosolic acetyl-CoA for fatty acid biosynthesis, is indispensable for cancer stemness. Overexpression of *CCND1* rescues the phenotype of *ITGA2* knockdown-induced cell cycle arrest. High expression levels of the *ITGA2*/*ACLY/CCND1* axis are correlated with an unfavorable relapse-free survival of patients with high grade breast cancer, in both basal-like and other subtypes. This study identifies *ITGA2* as a potential therapeutic target of TNBC stemness and metastasis.

## Introduction

Advancement of cancer medicine demands better prevention and blockage of cancer dissemination and metastatic disease that account for ∼90% of cancer related deaths, estimated 9.6 million globally in 2018^1^. Breast cancer is the most common cancer in women other than skin cancer and its five-year survival rate decreases to 27% when distant metastases develop ^2, 3^. Among all breast cancers, one of the most aggressive subtypes with early metastases is triple negative breast cancer (TNBC) which has the most limited options for targeted therapy and is associated with a poor prognosis ^4^. Therefore, it is imperative to identify the metastasis drivers and novel targets of TNBC progression.

Over the last two decades, human tumor cells with stem cell properties (stemness) have been identified and demonstrated to initiate tumorigenesis and accelerate progression, coupled with therapy resistance, immune evasion and distant metastases ^5–12^. However, targeting cancer stemness has yet to be widely achieved in the cancer clinic and therefore demands a better understanding and identification of targetable stemness regulators. While genetic and epigenetic aberrations are known to enable tumor plasticity and stemness ^10, 13^, we focuses on epigenetic regulation in cross talk with micro-environmental cues in this work.

Our previous studies have identified a few microRNAs (miRs) that regulate breast cancer stem cell-mediated chemoresistance and progression in TNBC ^14–18^. Specifically, miR-206 suppresses breast cancer stemness and lung metastasis ^14^. Comprising a class of small non-coding RNA sequences of 20-22 nucleotides, miRs mediate posttranscriptional regulation of target genes. Upon binding to the partial complementary sites of selective mRNAs, which are mostly located at the 3’ untranslated regions (UTR), miRs inhibit translation or accelerate degradation of their target mRNAs ^19–21^. Microarray-based global transcriptome analyses revealed that *ITGA2,* which encodes a surface integrin receptor CD49b (integrin α2), is one of the genes suppressed by miR-206.

Integrins include a family of adhesion molecules with binding affinity for various extracellular membrane components. Many integrins are key players in physiological and pathological processes in αβ heterodimers ^22^. CD49b is one of the integrin α subunits forming heterodimers with the β1 integrin subunit (CD29) and can be specifically activated by binding to collagen through its extracellular region ^23^. CD49b is known to be expressed in immune cells especially in regulatory T cells (Treg) ^24^. However, its intrinsic role and regulatory signaling in breast cancer stemness and progression have yet to be fully elucidated.

In this study, we utilized multiple TNBC model systems *in vitro* and *in vivo* to show that CD49b enhances breast cancer stemness and metastasis. Clinical correlation studies further demonstrate that *ITGA2* expression coincides with breast cancer outcomes.

## Results

### *ITGA2* is a target of miR-206 enhancing mammosphere formation

Our previous work revealed that miR-206 suppresses breast cancer stemness and metastasis ^14^. Based on the microarray dataset GEO-GSE 59751, we found that CD49b-encoding *ITGA2* mRNA levels were specifically decreased by miR-206 whereas other integrin genes such as *ITGB1* levels remained unaffected (**Supplementary Fig S1A**). Compared to the significant lung metastases mediated by control cells, the miR-206 expressing TNBC cells caused minimal, residual lung metastases with reduced CD49b levels upon tail vein infusion (**Supplementary Fig S1B**). The inhibitory effects of miR-206 on *ITGA2* mRNA and CD49b protein expression (both cellular and surface protein levels) were further validated in MDA-MB-231 cells upon transient transfection of miR-206 mimics, measured by qRT-PCR, immunoblotting, and flow cytometry analyses (**Fig 1A-C**, **Supplementary Fig S1A**).

**Figure 1.**
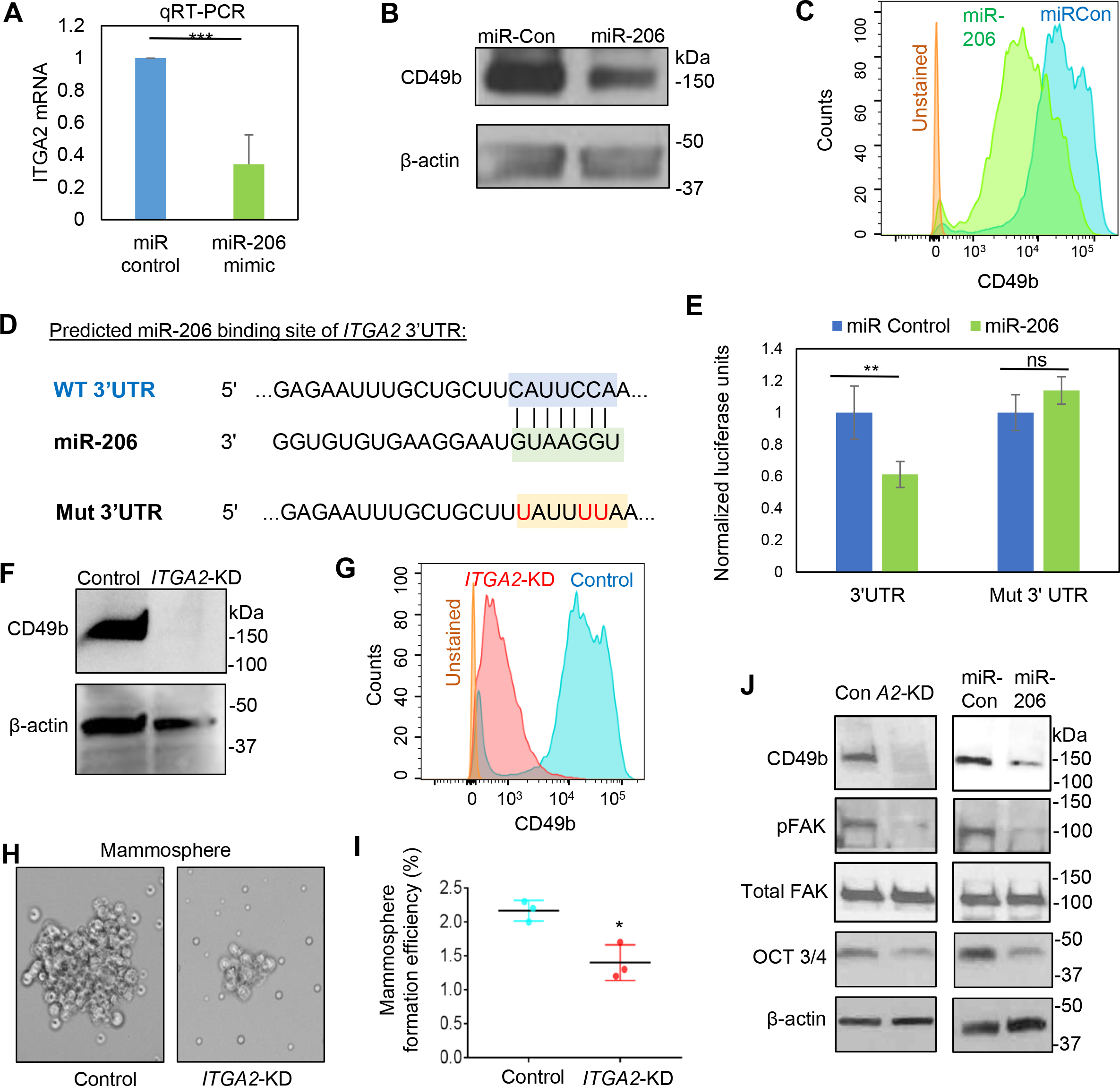
*ITGA2* is a target of miR-206 that promotes mammosphere formation and stemness factor expression. **A.** *ITGA2* mRNA levels are reduced by miR-206 transfection in MDA-MB-231 cells, measured via q-PCR. **B** & **C.** miR-206 inhibits *ITGA2*-encoded CD49b protein expression in MDA-MB-231 cells, evaluated via immunoblotting (B) and flow cytometry analyses (C). **D.** Predicted binding sites between the complementary sequences of *ITGA2* 3’UTR (WT) and miR-206. The bottom line shows the mutated C to T (U, in red) within the interaction site of 3’UTR. **E.** 3’UTR luciferase assay shows direct regulation of *ITGA2* by miR-206, n=3, T test p**=0.005. **F** & **G.** siITGA2 transfection depletes CD49b protein expression, measured by immunoblotting (F) and flow cytometry analyses (G). **H** & **I.** Images (H) and quantified counts (I) of breast cancer cell-derived mammospheres upon si*ITGA2* knockdown. **J.** Immunoblots to detect reduced expression of CD49b, phosphorylated FAK (pFAK), and OCT3/4 levels without affecting total FAK levels upon *ITGA2* KD or miR-206 upregulation.

To determine if *ITGA2* is a direct target of miR-206, we utilized multiple algorithms including TARGET SCAN and others to assess the *ITGA2* sequence and identified a predicted miR-206 binding site within its 3’UTR region (**Fig 1D**). The *ITGA2* 3’UTR region (2kb) was then cloned downstream of a luciferase reporter gene in pLS_3UTR. The luciferase reporter expression/activity was inhibited by co-transfected miR-206 and the inhibition was reversed when the three nucleotides were mutated from C to T/U within the predicted binding site for miR-206 (**Fig 1D-E**). These data suggest that *ITGA2* is inhibited by miR-206 through a direct binding to its 3’UTR site.

We can continued to examine the importance of *ITGA2* in regulating cancer stemness by knocking down its expression in MDA-MB-231 cells, and their effectiveness was confirmed by both immunoblotting and flow analyses (**Fig 1F-G**). Mimicking the mammosphere-inhibitory phenotype of miR-206 as we previously observed ^14^, *ITGA2* knockdown (KD) suppressed mammosphere formation of MDA-MB-231 cancer cells (**Fig 1H-I**). Furthermore, we investigated the effects of miR-206 upregulation and *ITGA2* KD on pluripotency transcription factors and focal adhesion kinase (FAK) signaling. FAK is known to be activated by many integrins ^25^ and the FAK signaling pathway promotes stemness in breast cancer cells ^26^. Indeed, miR-206 upregulation and *ITGA2* KD similarly decreased the protein levels of pluripotency marker OCT 3/4 and both inhibited the phosphorylation/activation of FAK, without altering the total FAK protein levels (**Fig 1J**). These results demonstrate that ITGA2/CD49b is intrinsically required to enhance cancer stemness in TNBC.

### *ITGA2* knockdown reduces cell cycle and proliferation

We further investigated whether CD49b regulates cancer stemness by modulating cell cycle or cellular viability. Similar to the miR-206-induced effects, *ITGA2* KD reduced confluency (**Fig 2A, Supplementary Fig S2A-C**) and counts of MDA-MB-231 breast cancer cells (**Fig2B**). We then examined if alterations in cell cycle and cell death contribute to the phenotypic changes. Mimicking the effects of miR-206 upregulation, *ITGA2* KD in MDA-MB-231 cells increased the percentage of cells in the G1 phase and decreased populations in cycling phases such as G2/M (**Fig 2C-D**). Consistently, in a different TNBC line BT-20, *ITGA2* KD also resulted in an accumulation of G1 arrested cells (**Fig 2E, Supplementary Fig S2C-D**). As expected, overexpression of CD49b caused an opposite effect in MDA-MB-231 cells and increased the percentage of cells in G2/M (**Fig 2F**).

**Figure 2.**
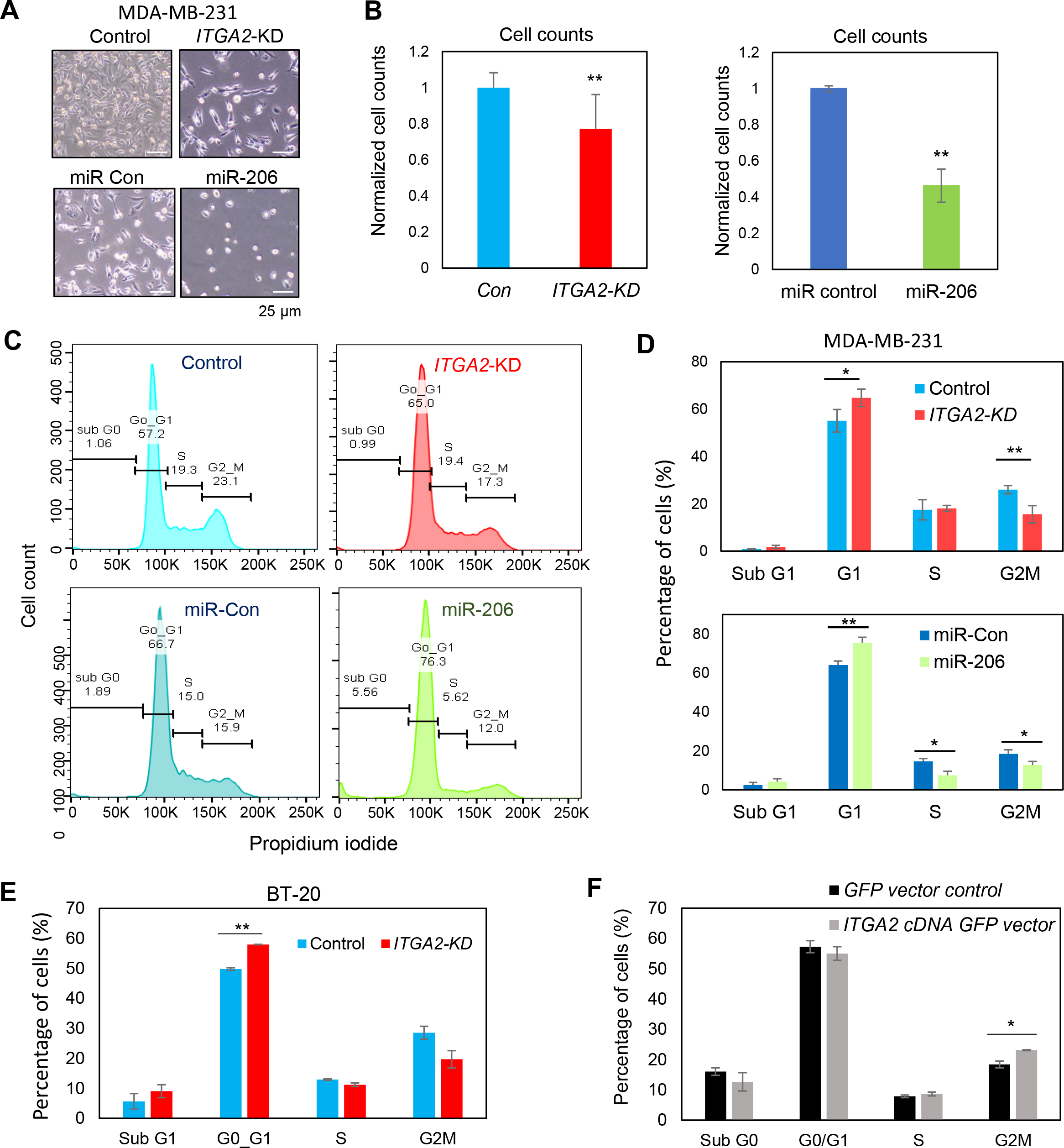
*ITGA2* knockdown inhibits cell cycle. **A** & **B**. Images (A) and cell counts (B) of MDA-MB-231 cells 48 hours after transfections with si*ITGA2* (*ITGA2*-KD) or miR-206 mimic. Cell number was measured via hemocytometer counting (B), n=12, p=0.002 (*ITGA2*-KD), and n=3, p=0.001231 (miR- 206). **C.** Representative flow cytometry analyses of cell cycle with phases of subG1, G1, S, and G2M using propidium iodide upon *ITGA2*-KD and miR-206 overexpression in MDA- MB-231 cells. **D.** Quantified percentage of MDA-MB-231 cells in each cell cycle phase as shown in C. Top panel: si*ITGA2* mediated *ITGA2-*KD and control, n=4 biological replicates; p=0.03 (G1); p=0.004 (G2M). Bottom panel: miR-206 overexpression and control, n=3 biological replicates; p=0.01 (G1), 0.02 (S), 0.04 (G2M). **E.** Percentage of BT20 cells in each cell cycle phase upon *ITGA2*-KD. **F.** Percentage of MDA-MB-231 cells in each cell cycle phase upon *ITGA2* cDNA- mediated overexpression (gated GFP^+^ cells with *ITGA2* overexpression).

We further measured cell death in breast cancer cells upon altering CD49b expression. From the cell cycle analyses, *ITGA2* KD did not cause significant changes of subG1 phase (**Fig 2C-D**), and differences in levels of cell death were not detectable in either Annexin V or propidium iodide (PI)-stained cells (**Supplementary Fig S3A-B**). A DNA binding dye with red fluorescence only marked a marginal increase of cytotoxicity (from 2% to 5%) of *ITGA2* KD cells during cell migration in a scratch wound healing assay (**Supplementary Fig S3C-D**). Furthermore, in order to ensure minimal off-target effects of siRNAs, we employed four individual *ITGA2* siRNAs #1-4 for additional validation studies. Most of the si*ITGA2s* had minimal effects on cell death, measured by Annexin V and PI-staining, as well as subG1 populations (**Supplementary Fig 3E-G**). Therefore, cell death was excluded as a major factor driving cell count differences which are likely to be primarily caused by cell cycle arrest. These data demonstrate that *ITGA2* promotes cell cycle which is required for proliferation and self-renewal of TNBC.

### *ITGA2* knockdown inhibits tumor cell migration and invasion

We subsequently investigated the functions of *ITGA2/*CD49b in regulating metastasis-related phenotypes, such as migration and invasion in multiple human and mouse TNBC cell lines. Collagen I is one of the primary ligands of integrins, which can bind extracellularly through the MIDAS motif and activate the integrin signaling cascade ^23^. To assess the impact of CD49b suppression on Collagen I binding, migration and invasion assays were conducted on collagen I-coated dishes. Transfection of a smart pool of 4 si*ITGA2*s significantly suppressed migration of MDA-MB-231 cells (**Fig 3A-C**). In addition, the siRNAs that conveyed the most effective reduction of CD49 levels as measured by flow cytometry also led to the most dramatic decrease in cell density and cell migration (**Supplementary Fig 3A-B**). Subsequently, cell invasion was measured when migrating cells were overlaid with Matrigel after the wound scratch was made. Upon *ITGA2* KD, MDA-MB-231 cells exhibited a dramatic decrease of invasion along with reduced cell density over time compared to the controls (**Fig 3D-F**). Cell migration was also inhibited in mouse TNBC 4T1 cells when transfected with siRNAs targeting mouse *Itga2* (**Fig 3G-I**).

**Figure 3.**
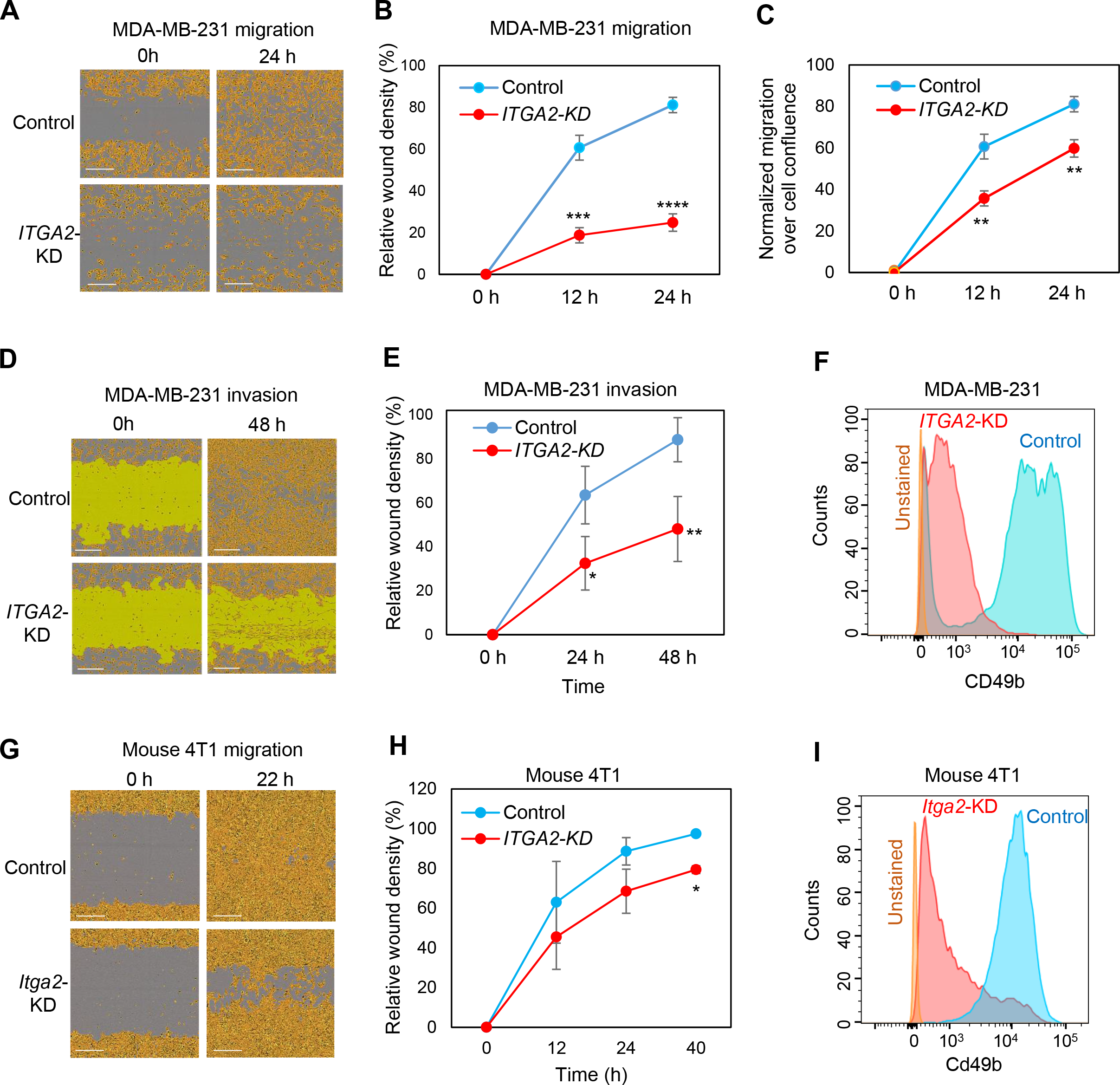
*ITGA2* knockdown reduces migration and invasion of TNBC cells in the presence of collagen. **A.** Images of migratory MDA-MB-231 cells with human siITGA2 smart pool-mediated knockdown (*ITGA2*-KD) and siRNA controls (Con), at 0 and 24 hour (h) following scratched wounding. Cells were plated on collagen I-coated plates. **B.** Quantification of relative cell density filled by *ITGA2*-KD and control cells in wounded areas, n=3; p=0.0002 (12 h), 0.0001 (24 h). **C.** Normalized migration over confluence of MDA-MB-231 cells during wound healing. **D** & **E**. Images (D) and relative cell density (E) of invasive MDA-MB-231 cells (A2-KD and control) covered by Matrigel, at 0 and 48 h after scratched wounding. n=3, p=0.02 (24 h), 0.007 (48 h). **F.** Flow cytometry plots of CD49b levels in MDA-MD-231 cells upon *ITGA2* KD. **G-H.** Images (G) and quantified wound density (H) of murine 4T1 mammary cancer cells upon *Itga2* KD via si*Itga2*, within 12, 22 (24), and 40 hours following scratch wounding. *t-test p<0.05 (n=3). **I.** Flow cytometry analyses of reduced murine Cd49b levels upon *Itga2* KD in 4T1 cells.

Supporting the potential for CD49b serving as a new therapeutic target to block cancer progression, we further found that an anti-CD49b neutralizing antibody blocked the migration of MDA-MB-231 cells during wound healing (**Supplementary Fig S4C**).

### *ITGA2* knockdown blocks lung colonization and metastasis

To examine the importance of *ITGA2* gene in TNBC metastasis, we set out to determine if *ITGA2* knockdown (KD) mimics miR-206 overexpression in regulating mammary cancer metastasis *in vivo*. Stable selection of sh*ITGA2*-transduced cells could not be achieved due to noted cell cycle defects. Thus, we focused on using transient gene silencing within the feasible period of time to perform lung colonization and spontaneous metastasis studies with human MDA-MB-231 and murine 4T1 cells, respectively. Upon tail vein injection, MDA-MB-231 cells with reduced CD49b levels had significantly decreased lung outgrowths in NOD/SCID mice (**Fig 4A-C**). While equivalent cells of the control and *ITGA2-*KD groups homed to the lungs immediately after tail vein injection, cells with *ITGA2-*KD had a reduced capacity to colonize the lungs as measured from 24 hour to 4 days post injection.

**Figure 4.**
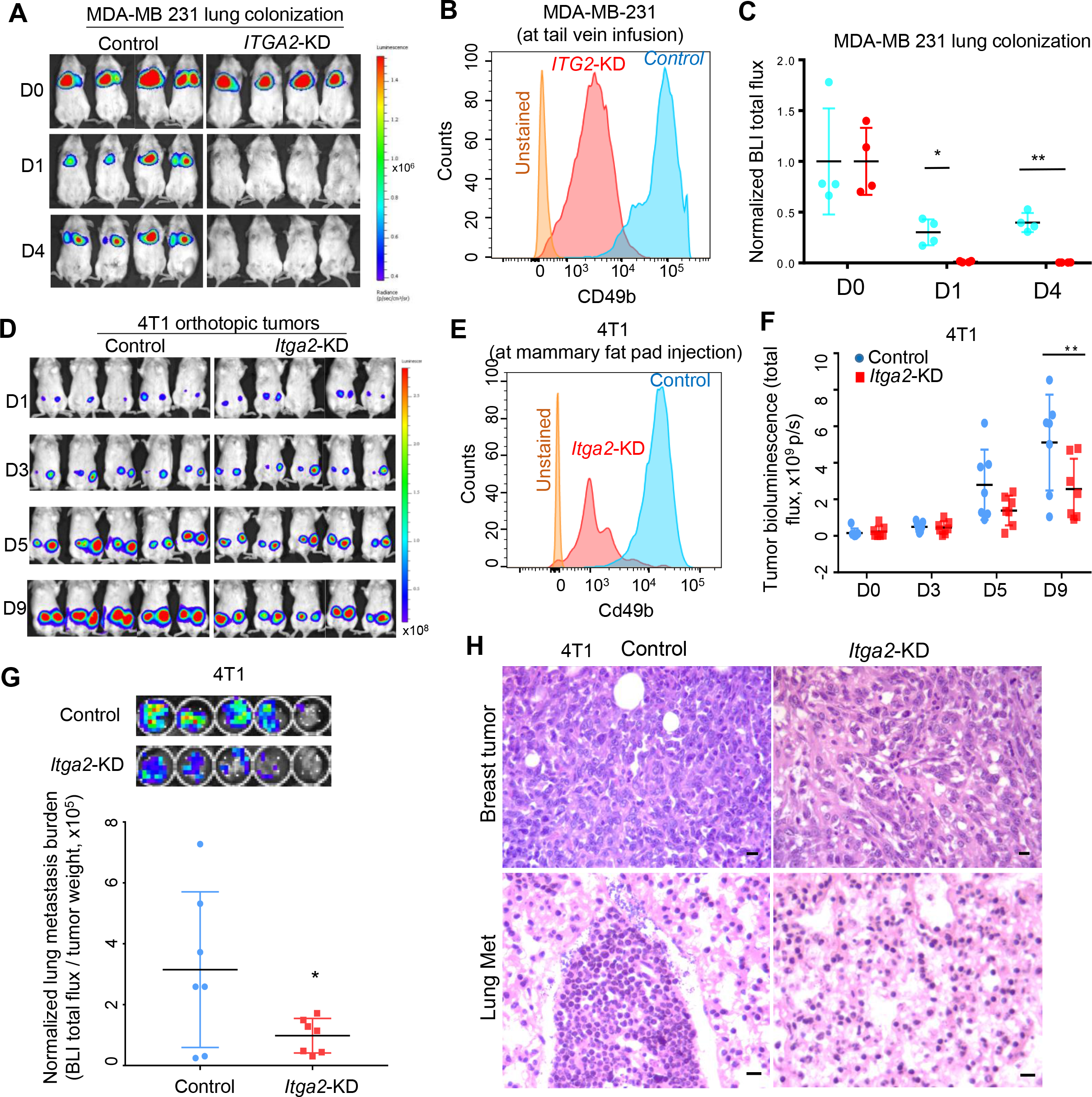
*ITGA2* knockdown inhibits TNBC lung metastasis. **A** & **B.** Bioluminescence images (A) and normalized lung colonization signals (B, ratio of the total flux compared to D0) of NOD/Scid mice on day 0 (D0), 1 (D1), and 4 (D4) post tail vein infusion of L2T/L2G-labeled MDA-MB-231 cells, control and *ITGA2*- KD. n=4 p=0.02 (D1) and 0.004 (D4) for paired comparisons. **C.** Flow analyses of CD49b in cells with or without *ITGA2*-KD, prior to tail vein infusion. **D** & **E.** Bioluminescence images (A) and tumor growth signals (B, total flux, n=7, p< 0.01) of mouse 4T1 cells, control and *Itga2* KD, on day 1 (D1), 3 (D3), 5 (D5), and 9 (D9) post orthotopic implantation into the 4^th^ mammary fat pads of BALB/c mice. **F.** Flow analyses of mouse Cd49b levels in 4T1 cells +/- *Itga2*-KD prior to implantation. **G.** Bioluminescence images (top panels) and normalized lung metastasis by primary tumor burden (total flux/tumor weight) of the mouse lungs bearing 4T1 spontaneous metastases on D9, n=7, p=0.05. **H.** H & E staining images of 4T1 breast tumors and spontaneous lung metastases (present in the control group and absent in the *Itga2* KD group) on day 9 post orthotopic implantations shown in D. Scale bars = 20 µm.

In addition, the murine 4T1 TNBC cells were utilized to investigate the effects of *Itga2* KD on spontaneous metastasis in immune-competent Balb/c mice. When these cells were orthotopically transplanted at large numbers (10^6^ cells per injection), 4T1 cells could metastasize from the mammary fat pads to the lungs within a week ^27^. Using IVIS imaging of luciferase-labeled cells, we found that silencing *Itga2* gene and Cd49b expression only modestly compromised primary tumor growth (**Fig 4D-F**), but dramatically suppressed lung metastasis (normalized on tumor weight) (**Fig 4G, Supplementary Fig S1C-D**). Reduced lung metastases of *Itga2-KD* 4T1 cells was validated by H & E staining (**Fig 4H**). In contrast to the lungs, metastasis of 4T1 to the liver and other organs was not detected in these mice within the short observation window.

### *ITGA2* regulates lipid metabolism and cell cycle pathways

To further understand the downstream pathways regulated by *ITGA2* in breast cancer, we performed RNA sequencing to compare the transcriptomes of *ITGA2* KD and control MDA-MB-231 cells and identified the top 195 differentially expressed genes, based on a cutoff of FDR < 0.05 and Log2 (Fold Change) > 0.48 or < −0.48 (**Fig 5A**). Notably, based on Metascape analysis (http://metascape.org), *ITGA2* KD altered the pathways of lipid metabolism, retinoid metabolism and transport, cell cycle, response to wounding, regulation of protein kinase activity, and other pathways, including representative genes such as *ACLY, CCND1, CCND3, MCM5, TGFB2, PCNA, ADAMTS1, MAP3K5* (**Fig 5B-E, Supplementary Fig S5**). ACLY-mediated acetyl-CoA production for lipid metabolism was recently shown to promote tumorigenesis in pancreatic cancer ^28^. *CCND1* appeared in multiple pathways regulated by *ITGA2*/CD49b, such as cell cycle, response to stress (toxic substance), regulation of protein kinase activity (serine/tyrosine phosphorylations), cytokine signaling, unfolded protein response, mitotic cell phase transition, androgen receptor (AR) pathway, etc (**Fig 5B, Supplementary Fig S5**). We chose ACLY and CCND1 for further functional studies to determine their importance in CD49b-mediated phenotypes in TNBC.

**Figure 5.**
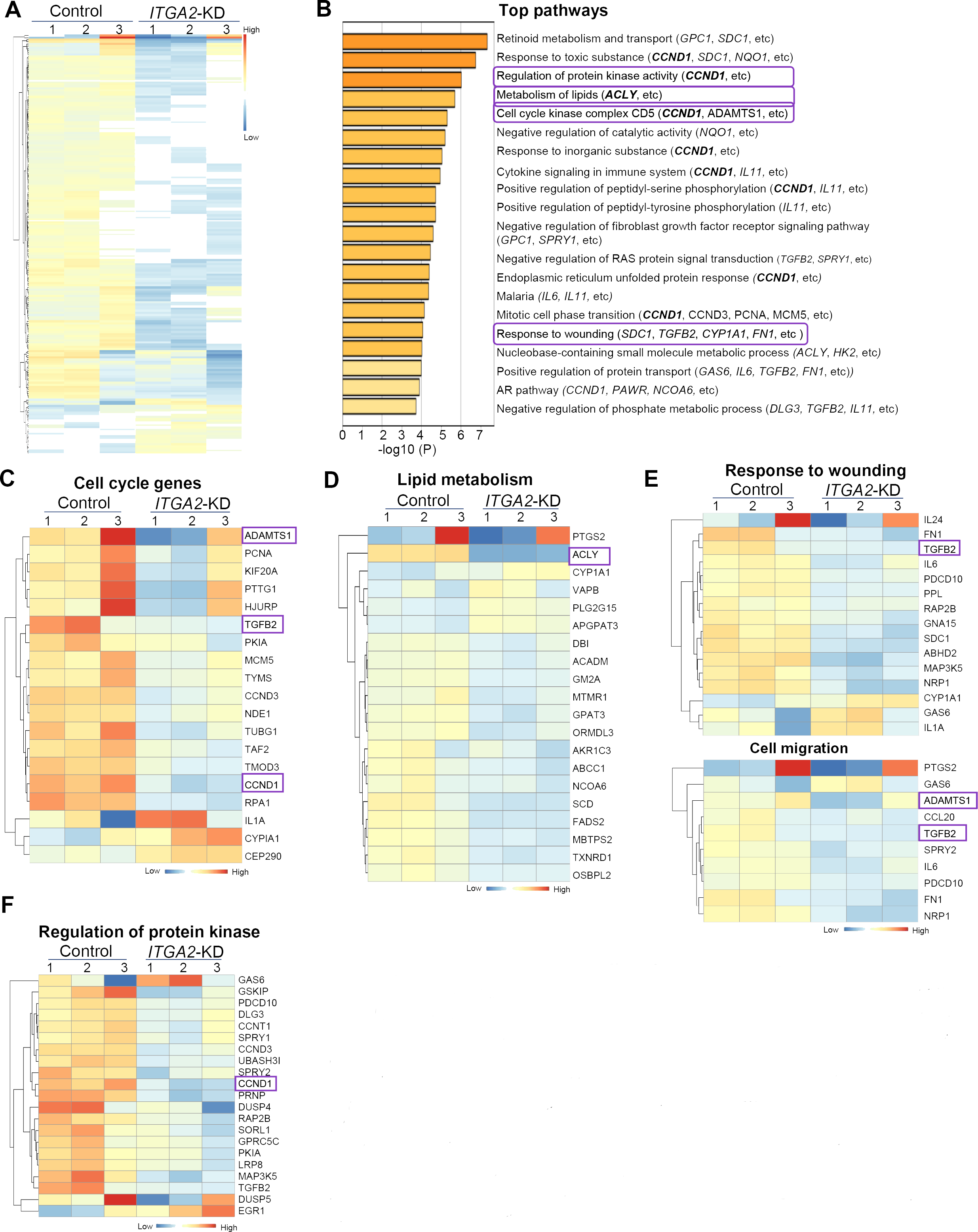
*ITGA2* downregulation-altered genes and clusters. **A.** Heatmap of differentially expressed 195 genes >1.5 fold change upon *ITGA2* KD via si*ITGA2* (see Supplementary Table S1), with most being downregulated (top and middle rows of 169 genes) and a small portion of upregulated genes (bottom rows of 26 genes). **B.** Top 20 gene clusters enriched by *ITGA2* KD, colored by p-values. The Metascape gene list analyses were carried out with the ontology sources of KEGG Pathway, GO, Reactome, Canonical Pathways, and CORUM. **C-F.** Heatmap of genes in representative pathways of cell cycle, lipid metabolism, wound healing and cell migration, and regulation of protein kinase activity, with a few genes shown in multiple pathways possibly regulating self-renewal and lipid metabolism, such as *CCND1* and *ACLY*.

### ACLY and Cyclin D1 are CD49b targets enhancing cancer stemness and cell cycling

ACLY was one of the top gene hits downregulated in *ITGA2* KD cells and hypothesized as a regulator of cancer stemness in breast cancer. We first confirmed decreased protein levels of ACLY in *ITGA2* KD cells as well as *ACLY* KD cells (**Fig 6A**). *ACLY* KD indeed mimicked *ITGA2* KD in inhibiting mammosphere formation (**Fig 6B**), without significantly altering total cell counts or cell motility (**Supplementary Fig S6**), suggesting a role of ACLY in stemness regulation independent cell cycle regulation.

**Figure 6.**
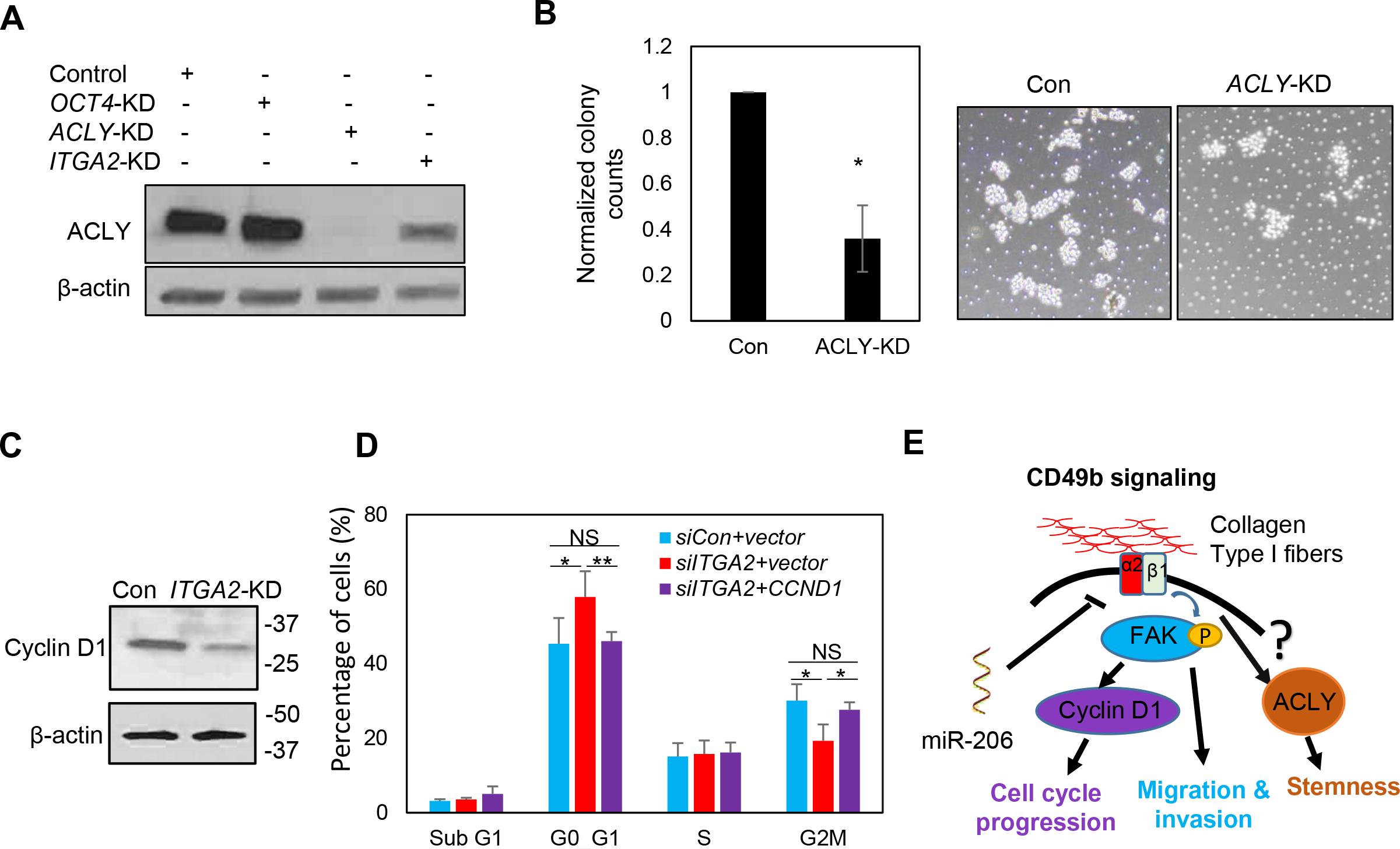
ACLY and Cyclin D1 are CD49b targets enhancing self-renewal and cell cycle. **A.** Immunoblot of downregulated ACLY proteins in MDA-MB-231 cells transfected with si*ACLY* and si*ITGA2* in comparison to siRNA control and si*OCT4*. **B.** Reduced mammosphere formation in *ACLY* KD cells, shown in quantified histograms (left panel) (p<0.05, n=3 biological replicates) and representative images (right panels). **C.** Immunoblot of downregulated cyclin D1 proteins in MDA-MB-231 cells transfected with si*ITGA2* compared to control. **D.** si*ITGA2*-induced G1 arrest and altered G2/M phase were rescued by CCND1 overexpression. **E.** Schematic CD49b signaling pathway induced by extracellular matrix factors such as collagen I fibers that result in the phosphorylation of FAK and upregulation of *CCND1* levels to promote proliferation of stem/progenitor cells and metastasis of TNBC.

We then continued to determine if CD49b-regulated cell cycle attributes to *CCND1*. *ITGA2* KD also decreased the protein levels of *CCND1*-encoded cyclin D1 (**Fig 6C**), which is an important driver of the G1-S phase transition during the cell cycle ^29^. Furthermore, enforced expression of *CCND1* rescued the cell cycle defects in ITGA2 KD cells (**Fig 6D**). Meta-analysis of several breast cancer gene expression datasets using bc-GenExMiner further demonstrated a positive correlation of *ITGA2* with both *CCND1* and *ITGB1* levels, without a significant correlation between *CCND1* and *ITGAB1* levels (**Supplementary Fig S6**). Since FAK is known to promote *CCND1* expression, our data suggest that miR-206 inhibits *ITGA2*/CD49b-mediated signaling cascade through compromised FAK phosphorylation and cyclin D1 expression as well as other stemness and metastasis-regulating pathways (**Fig 6E**).

### *ITGA2* pathway components are associated with poor survival of breast cancer patients

Lastly, we examined the pathological and clinical relevance of *ITGA2* expression in breast cancer. Based on the gene expression profiles of over 50 established breast cancer cell lines, high *ITGA2* expression levels are mainly observed in basal A (BaA) subtype cells (**Supplementary Fig S7**). Relapse free survival (RFS) plots show that high expression levels of *ITGA2* are associated with reduced RFS probability in patients with the intrinsic basal subtype of breast cancer (**Fig 7A**), and all grade 3 breast cancer patients (**Fig 7B**) independent of ER expression (**Fig 7C-D**). Consistently, the analysis of another database also demonstrated the correlation of high *ITGA2* expression with poor event free survival of the patients with TNBC (**Fig 7E**). Similar to *ITGA2*-based KM Plots, *CCND1* and *ACLY* expression show a similar pattern of correlation with a short RFS in breast cancer patients (**Fig 7F-G**), whereas miR-206 expression indicates an extended RFS in ER^−^ breast cancer (**Fig 7H**). Our recent discoveries and other reports demonstrated that circulating tumor cell (CTC) clusters are enriched with cancer stemness and proliferation compared to single CTCs ^11, 30–32^. Notably, shown by single cell sequencing, both *ACLY* and *CCND1* expression levels are upregulated in CTC clusters versus single CTCs isolated from patients with breast cancer and patient-derived xenografts (**Fig 7I**), correlating with their effects on improved self-renewal and cell cycle progression respectively.

**Figure 7.**
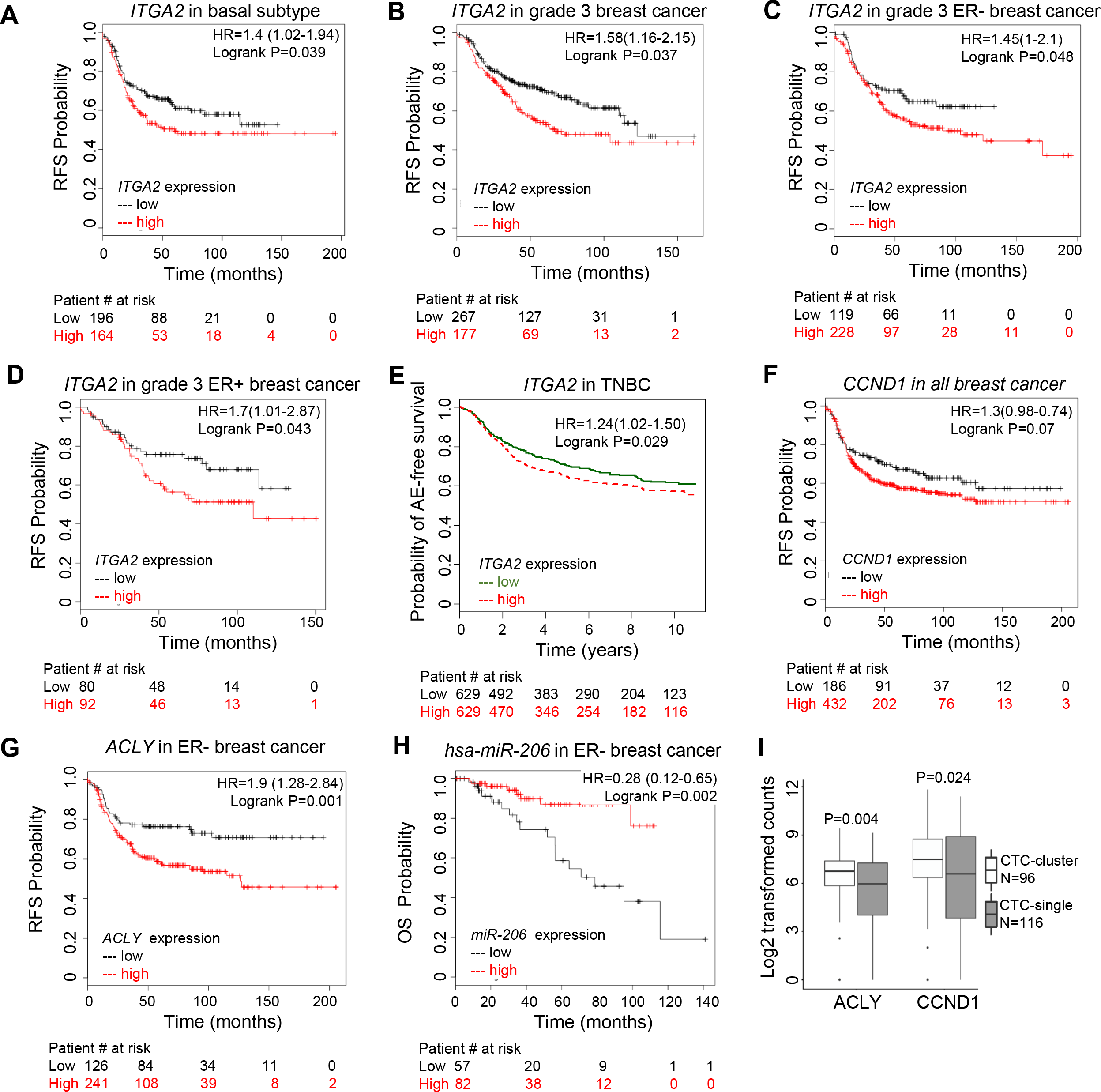
*ITGA2* regulation pathway is associated with survival of patients with breast cancer. **A.** KM plots of relapse free survival (RFS) associated with *ITGA2* expression in basal subtype of breast tumors, analyzed using Breast Cancer Kaplan-Meier Plotter, best cut off 46% of high expressing patients and 54% of low expressing cells, n=360 total patients; p=0.039. **B-D** KM plots of RFS correlation with *ITGA2* expression in all grade 3 (B, p=0.0037, n=444, cut off 40% high), grade 3 ER- (C, p= 0.048, n=347, cut off 66% high), and grade 3 ER+ (D, p=0.043, n=172, cut off 53% high) breast cancer, best cut off selected in Breast Cancer Kaplan-Meier Plotter analyses. E. Breast cancer miner plot of any-event (AE) free survival association with *ITGA2* expression in TNBC. p=0.02, n=1258, median cut off. F-H. KM plots of RFS or OS correlated with *CCND1* expression (F) in all breast cancer (p=0.07, n=618), *ACLY* expression (G) in ER- breast cancer (p=0.001, n=407), and *miR-206* expression (H) in ER- breast cancer (p=0.002, n=159), beast cut off. I. Box plot for differential *ACLY* and *CCND1* expression levels between clustered and single CTCs of breast cancer patients and PDXs.

Taken together, our studies identified the *miR-206*/*ITGA2/ACLY-CCND1* signaling cascade as a clinically relevant target for inhibiting breast cancer metastasis and potentially improving patient outcomes.

## Discussion

Our findings have identified that *ITGA2*/CD49b as a direct target of miR-206 in promoting breast cancer stemness and metastasis. The surface protein and integrin family member CD49b can be targeted by siRNAs and neutralizing antibodies, serving as a new therapeutic target for metastatic TNBC. FAK, a non-receptor tyrosine kinase is activated by CD49b signaling and has been identified as a key mediator of intracellular signaling by integrins ^33, 34^. Several other integrins have also been identified as markers of cancer stem cells, such as CD49f (integrin α6), integrin α7, and CD104 (integrin-β4) ^35–37^. Both integrins and FAK can play crucial roles in the maintenance of breast cancer stem cells ^26, 38^. Future studies should determine if CD49b promotes expression of the pluripotency transcription factor OCT 3/4 through the FAK signaling pathway to regulate the breast cancer stemness phenotype ^38^.

While quiescent normal and malignant stem cells may be slow cycling, activated stem cells undergo proliferation to generate and replenish the progenies. CD49b appears to activate breast cancer stem and progenitor cells for promoted cell proliferation and progression. Our global transcriptome analyses have discovered that CD49b regulates cell cycle, wound healing, and protein kinase pathways related to self-renewal and migration. Cell cycle progression through G1 phase in mammalian cells is precisely regulated by integrin-mediated adhesion to the extracellular matrix growth and growth factor binding to receptor tyrosine kinases (RTKs) ^39, 40^. These two classes of receptors coordinately promote activation of the G1 phase cyclin-dependent kinases (CDKs), CDK4/6, and increasing expression of G1 phase cyclins, Cyclin D1/Cyclin E, which then induces genes that mediate S phase entry. Our results show that transient *ITGA2* knockdown inhibited TNBC proliferation by inducing G1 arrest. Previous studies have shown that integrin α5β1 regulates Cyclin D1 expression by sustaining mitogen-activated protein kinase (MAPK) activity ^41^. Another study shows that integrins interact with the adaptor Shc through the alpha subunit to activate the MAPK pathway in response to integrin ligation ^42^.

The expression of CD49b in immune cells especially in regulatory T cells (Treg) ^24^ needs to be taken into consideration when it serves as a potential target of cancer stemness. While many integrins promote migration, invasion and proliferation of cancer cells, it is unclear how CD49b in the stroma and immune microenvironment affects tumor development and metastasis. From Norma Ramirez’s studies, *Itga2* knockout mice develop accelerated metastasis of spontaneous Her2 + mouse tumors ^43^. However, in their model, both tumor cells and stroma cells lack the expression of Cd49b. Therefore it is unclear how much the tumor-intrinsic regulatory effects and extrinsic effect of *Itga2* deficiency contribute to the metastatic phenotype. Our studies support a cancer cell intrinsic integrin signaling pathway that contributes to the metastatic potential of these cells. Future studies with conditional *Itga2* knockout in mammary epithelial cells and immune cells respectively will assist clarifying the extent of impact of *Itga2* on tumor *versus* stromal cells in the metastatic process. Nevertheless, our studies have elucidated the intrinsic role of CD49b/Cd49b in TNBC, particularly in response to transient and acute depletion without the selective pressure of compensatory gene expression from other integrin subunits. These data suggest that development of cancer-specific and metastasis-blocking ITGA2 neutralizing antibodies should be pursued with the goal of follow-up translational studies.

Altered metabolism has been connected to cellular reprograming. In addition to cell cycle and stemness-related pathways, our studies have revealed that there are multiple metabolic pathways that are regulated by *ITGA2*, such as lipid metabolism with ACLY as a notable component. ACLY has recently been revealed to positively promote tumorigenesis and cell plasticity via elevated acetyl-CoA abundance and global histone acetylation ^44^. Our studies confirm a modulation of ACLY expression by CD49b signaling pathway and a role of ACLY in breast cancer self-renewal, revealing an unprecedented connection of integrin CD49b signaling to ACLY-mediated lipid metabolism and cancer stemness. Lastly, it will be important to determine if this is specific to CD49b or shared with other integrin subunits.

## Acknowledgements

We would like to thank Huiping Liu lab members particularly, Dhwani Patel, Nurmaa Dashzeveg, Wenjing Chen, and Natalia Brokate for assisting this manuscript. We are grateful to Dai Horiuchi for sharing cell lines as well as equipment with our laboratory; Dr. Thomas J. Hope laboratory for sharing the IVIS instrument to image immune competent animals; and Dr. Marc L. Mendillo laboratory for sharing the Incucyte imaging system. We appreciate the Animal Imaging Center as well as the Center for Advanced Microscopy and Nikon Imaging Center at Northwestern University for assisting analysis of IVIS data and for providing equipment; the NUSeq Core Facility for the RNA sequencing and the Bioinformatics Core for the analysis.

This manuscript has been partially supported by NIH/NCI grants R00CA160638 (H.L.), and Supplement for Diversity (V.A.) and R01CA213843 (R.A.K), American Cancer Society grant ACS127951-RSG-15-025-01-CSM (H.L.); the Susan G. Komen Foundation CCR15332826 (H.L.) and CCR18548501 (X.L); the Department of Defense W81XWH-16-1-0021 (H.L.); the Lynn Sage Cancer Research Foundation (X.L. and H. L.); Northwestern University’s Endocrinology Training Grant T32DK007169-39 (A.H.); and start-up funds from Case Western Reserve University and at Northwestern University (H.L.).

## Materials and Methods

### Animal studies

All mice used in this study were kept in specific pathogen-free facilities in the Animal Resources Center at Northwestern University and Case Western Reserve University. All animal procedures complied with the NIH Guidelines for the Care and Use of Laboratory Animals and were approved by the respective Institutional Animal Care and Use Committees. Animals were randomized by age and weight. The exclusion criterion of mice from experiments was sickness or conditions unrelated to tumors. Sample sizes were determined based on the results of preliminary experiments. NSG and NOD-Scid mice were used for tail vein injection experiments with MDA-MB-231 cells and 4T1 cells expressing luciferase2-fused eGFP (pFU-L2G) or tdTomato (pFU-L2T) ^8^. For spontaneous metastasis experiments the murine mouse cell lines (4T1) were injected in the mammary fat pad of BALB/c mice. Imaging of all in vivo experiments was performed using a Perkin Elmer IVIS.

### Cell culture

MDA-MB-231 and BT20 cells were cultured in DMEM high glucose, supplemented with 10% FBS and 1% Pen/step antibiotic. 4T1 cells were cultured in RPMI media supplemented with 10% FBS and 1%Pen/strep antibiotic. MDA-MB-231 cells were obtained from ATCC, and BT20 and 4T1 cells were originally from ATCC and expanded by Dr. Dai Horiuchi. Transcriptome and phenotypic analyses of these cells matched the published profiles.

### Cell transfections

Transient transfection of breast cancer cells with Dharmacon small interfering RNAs (siRNAs), or microRNA mimics was performed once or in tandem using Dharmafect transfection reagents (siRNA control:D-001810-01-05, ITGA2siRNA: L-004566-00-0005, CCND1: L-003210-00-0020; set of 4 ITGA2 siRNA: LU-004566-00-0005; miR control: CN-001000-01-05). Cells were at least 70% confluent at the moment of transfection, siRNA concentration is 100nM per transfection. Most phenotypic analyses were conducted 2 days after the last transfection. Mammosphere formation *in vitro* and metastasis assays *in vivo* were done on the following day after the second transfection. For rescue experiments, cells were co-transfected with GFP empty vector or GFP *ITGA2* cDNA in addition to either miR control or miR-206; alternatively OFP vector control, OFP-*CCND1* cDNA vector in addition to either siRNA control or si*ITGA2* (Sino Biological, CV025, CV026, HG13024-ACG, HG10905-ACR). Transfection of vectors were made in serum free media, using PolyJet (Signa Gen Laboratories, SL100688).

### Mammosphere assay

Cells were plated at a low density of 1,000 cells in suspension in a 6-well plate covered with poly-HEMA in PRIME-XV® Tumorsphere serum-free medium (Irvine scientific, 91130). 8 days after plating a total number of spheres (diameter >50 µm) were counted for each well and pictures were taken.

### Scratch wound assay

Cells were plated in an image lock 96 well plate overnight at a confluency of 20K cells per well. On next day morning a scratch was performed using the IncuCyte wound maker and cells were monitored by Incucyte. Plates were coated with collagen type I (Corning 354231) and incubated at RT for at least an hour prior to cell plating.

### Transwell invasion assay

24-well plate inserts were coated with Matrigel and cells were plated on the surface of the Matrigel (BD Sciences 356231) in serum free media. The bottom of the well were covered with media containing 10% FBS. Invasion of cells was measured after dissociation of the cells using a cell dissociation solution (Cultrex, 3455-096-05) from the insert and staining with calcein am (465 excitation, 535 for emission). Signal was measured using a BioTek fluorescence plate reader and normalized to control.

### Cell cycle analysis

Cells were collected, then fixed with 70% alcohol, washed with PBS, incubated with RNAse A for one hour then propidium iodide dye was added. Some cells were stained with DAPI and the RNAse A incubation was skipped. Cells were kept in the dark and at 4 degrees until flow cytometry analysis in LSRII (PE and UV channels).

### 3’UTR luciferase assay

Vectors expressing luciferase with the 3’UTR region of *ITGA2* as well as luciferase assay reagents were obtained through Active Motif and their protocol was followed (Switchgear, Lightswitch luciferase assay kit, 32031; GAPDH control, 32014; *ITGA2* 3’utr: pLS_3UTR).

### Flow cytometry

Cells were detached using 2mM EDTA in PBS and cell surface expression of CD49b was measured (Biolegend 359310). Annexin V staining was performed using FITC Annexin V apoptosis Detection Kit (BD Biosciences 556547).

### Western blot

Cells were lysed by RIPA buffer supplemented with Amresco protease inhibitor cocktail (1:100 diluted) and centrifuged for 10 mins at 4 degrees and 10,000 RPM. Protein concentration was measured through a Bradford assay and 20-40ug of protein were loaded for each sample. Antibodies used: Cd49b (Rabbit pAb, Thermo Fisher Scientific PA5-26061), pFAK, total FAK (Rabbit, Cell Signaling, mAb 8556 and pAB 3285), b-actin (Mouse mAb, Abcam ab8224). Secondary antibodies where HRP conjugated (Promega, Rabbit W401B, Mouse W402B) for ECL detection using Pierce ECL2 solution (Thermo Fisher Scientific, 1896433A).

### Immunohistochemistry

Tissue was first deparaffinized by xylene incubation followed by rehydration of tissue obtained through alcohol incubation in decreasing concentrations. Heat induced antigen retrieval was obtained using a decloacking chamber in decloacking solution for 20 mins (Biocare Medical, RD913L). Further staining of primary antibody was performed following Dako envision plus kit and DAB staining (Anti-*ITGA2*, Sigma HPA063556). Hematoxolin and eosin staining was performed following dehydration of the tissue by incubation with increasing concentrations of alcohol. Tissues were later mounted with permount.

### RNA sequencing analysis

Total RNA extraction was performed using Trizol and phase separation by chloroform, then extraction by alcohol. Samples were submitted to the RNA core for deep sequencing analysis. RNA sequencing was performed on a HiSeq 4000 at Northwestern University’s Center for Genetic Medicine Sequencing core facility and a library was made using TruSeq Total RNA-Seq Library Prep. Sequencing data was aligned using STAR ^45^ and analyzed using HTSeq ^46^ and DESeq2 ^47^. Differentially expressed genes were subject to a cutoff of FDR < 0.05 and Log2 (Fold Change) > 0.48 or < −0.48. Pathway analysis was performed on significantly-differentially expressed genes in Metascape (http://metascape.org) ^48^. The single cell RNA sequencing data of circulating tumor cells isolated from breast cancer patients and xenografts were retrieved from the published databases ^30, 32^ and compared using Wilcox T test.

### Human database and cell line expression analyses

Relapse free survival plots were obtained from KM-Plot (www.Kmplot.com/analysis) ^49^. This tool downloads gene expression data, relapse free and overall survival information from GEO (Affymetrix microarrays only), EGA and TCGA. It uses a PostgreSQL server, which integrates gene expression and clinical data simultaneously. KM-Plot analyzes the prognostic value of a particular gene by splitting the patient samples into two groups according to various quantile expressions of the proposed biomarker and the best cut off was chosen for each analysis. The hazard ratio with 95% confidence intervals and logrank P values were calculated.

Breast cancer miner was used for the analysis of any event free survival of basal like and/or TNBC (IHC-based) patients. This tool uses GSE studies as well others independent of KM plot ^50, 51^. The total amount of patients analyzed was 876 (filtered from 5,861).

UCSF Xena browser was used to study the ITGA2 gene expression through different breast cancer cell lines according to their subtype ^52^. The profiles of the set of breast cancer cell lines are seen in the study by Heiser 2012 ^53^.

### Statistical analysis

Student’s *t-*test was performed for most of the comparisons except for the ones specified such as in human database analyses. Probabilities under 0.05 were considered significant and represented with one star (*). Probabilities under 0.01 and 0.001 were represented with two stars (**) and three stars (***), respectively.

---

**Supplementary Figure S1.**
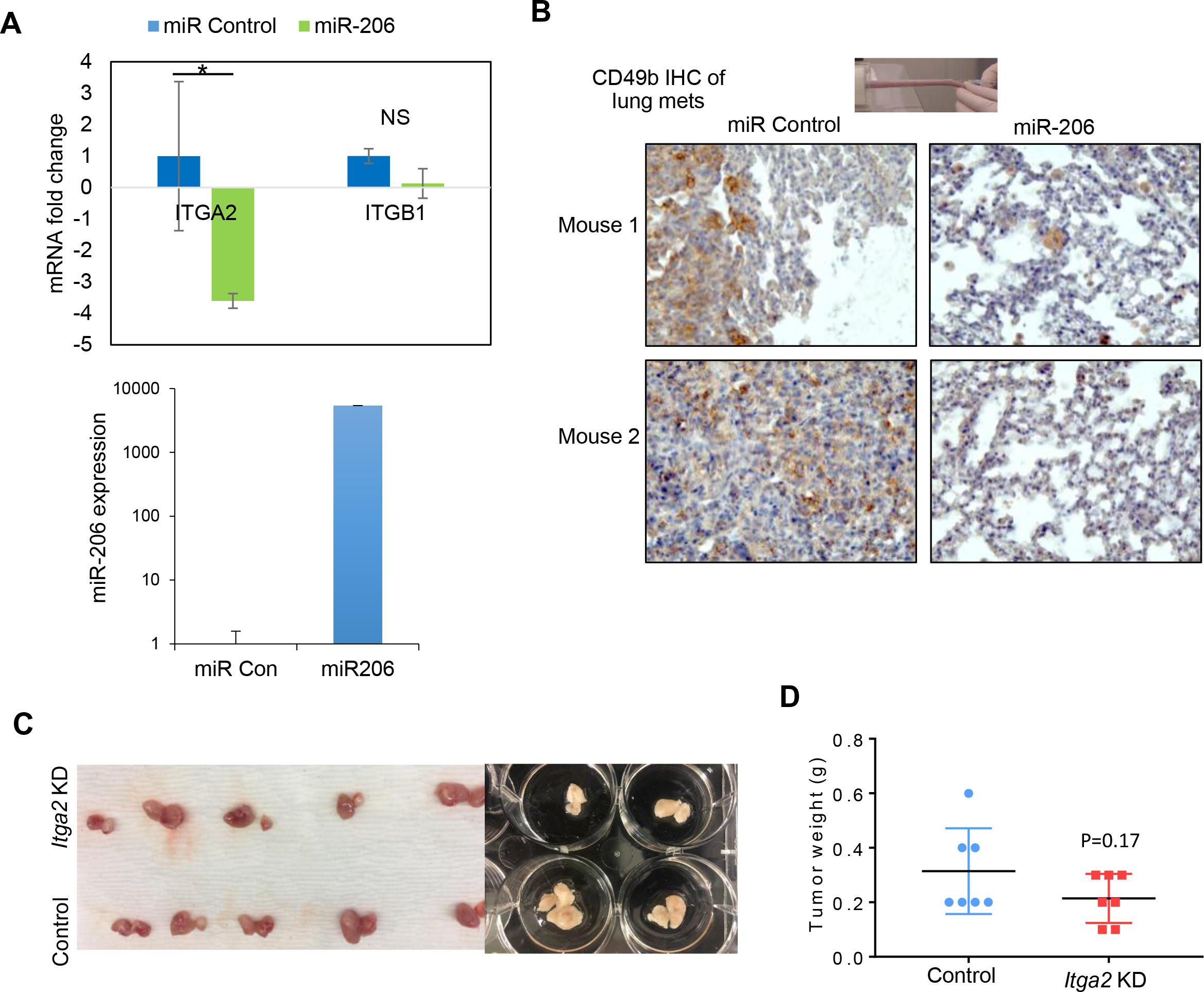
miR-206 inhibits *ITGA2* expression (related to Fig 1) A. Normalized microarray data showing reduced gene expression of ITGA2 (not ITGB1) by transient miR-206 overexpression. B. CD49b IHC staining of lung sections of mice 72 days post tail vein injection of TNBC MDA-MB-231 cells transfected with miR-206 and the miR control, respectively. C&D. The images (C) and weight (D) of 4T1 breast tumors mediated by cells transfected with siRNA control and siItga2 and orthotopically transplanted into the mammary fat pads of balb/c mice

**Supplementary Figure S2.**
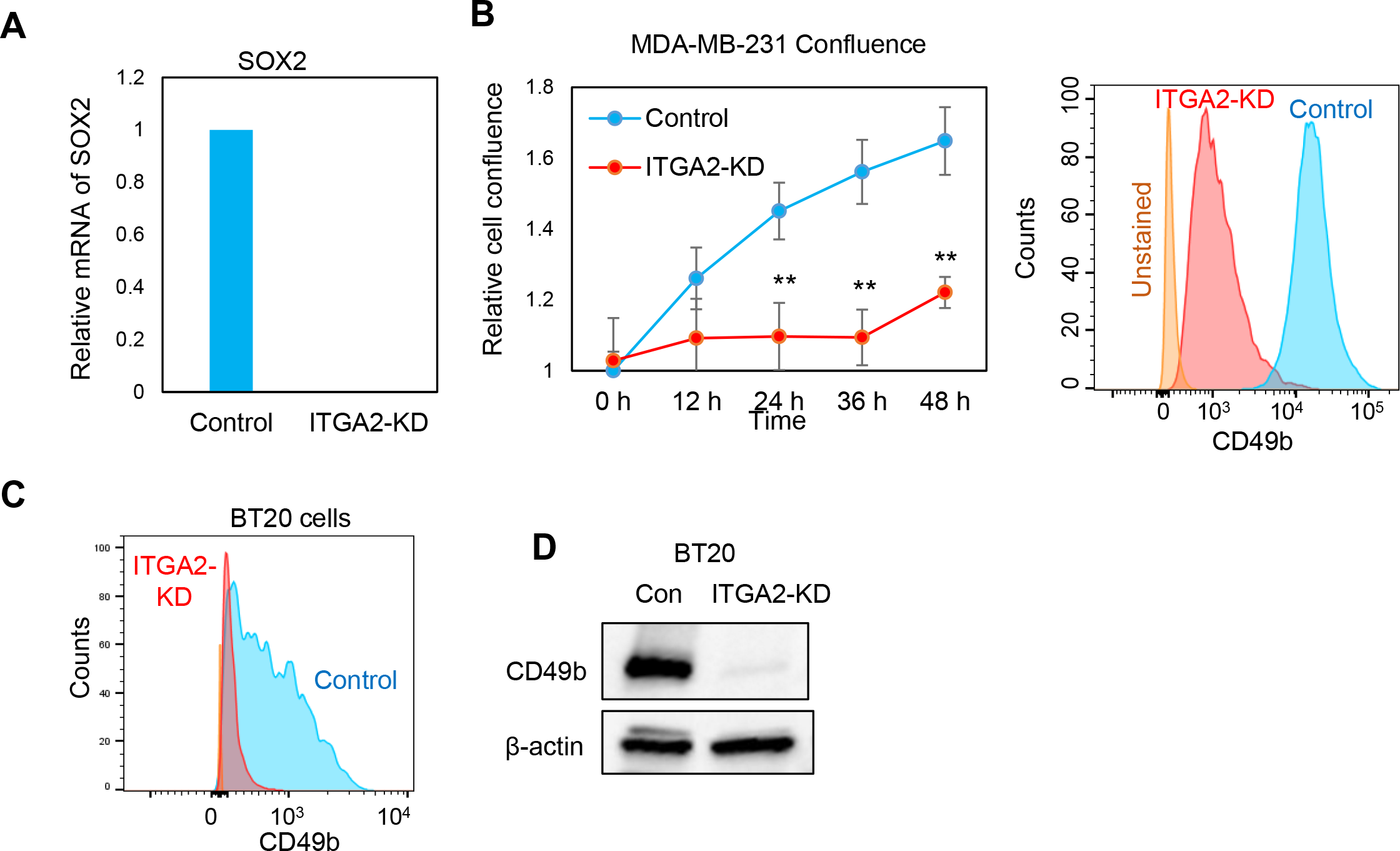
*ITGA2* KD reduces SOX2 expression and cell confluence (related to Fig 2) *A. ITGA2* KD decreased SOX2 mRNA levels in MDA-MB-231 cells. *B.* Confluence and CD49b expression of MDA-MB-231 cells upon *ITGA2* KD. **T test p<0.01. *C.* Reduced CD49b in BT20 cells upon *ITGA2* KD. *D.* Immunoblots for CD49b of BT20 cells upon *ITGA2* KD.

**Supplementary Fig S3.**
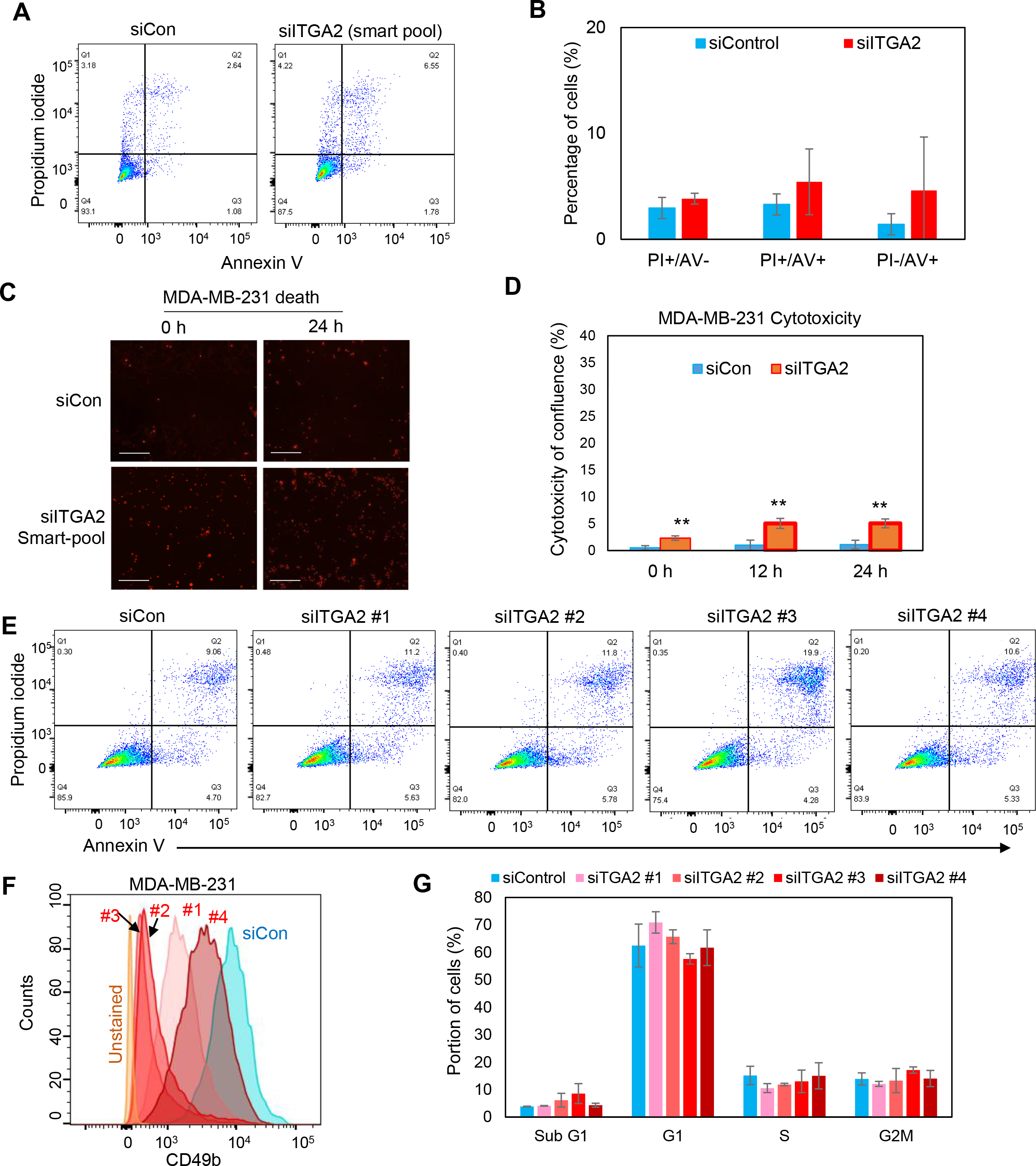
pooled and single si*ITGA2*s caused minimal death of MDA-MB-231 cells (related to Fig 3). **A**&**B**, Representative flow dot plots (A) and quantification (B) of cell death measure by Annexin V (AV) /Propidium Iodide (PI), PI+/AV+, n=2. **C**&**D,** siITGA2 Smart pool-induced cell death measured by cytotoxic red dye during wound healing (n=3, T test p**<0.01). **E.** Flow plots of AV^+^PI^+^ MDA-MB-231 cells transfected with individual siITGA2 #1-4. **F** & **G.** Flow analyses of CD49b (F) and cell cycle (G) in MDA-MB-231 cells transfected with individual siITGA2 #1-4.

**Supplementary Fig S4.**
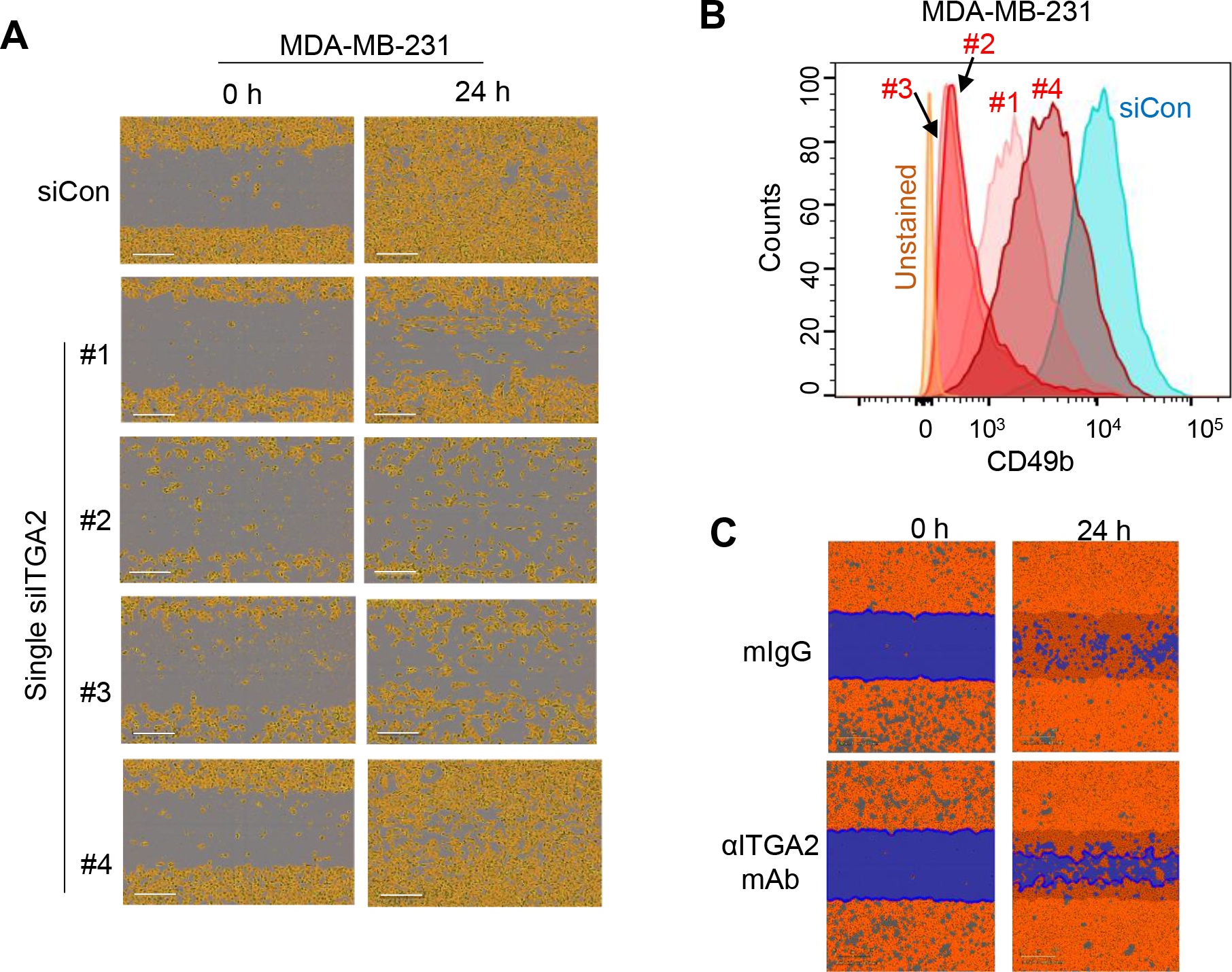
Effects of *ITGA2/Itga2* KD on cell migration and tumor growth (related to Fig 4). A. Migration of MDA-MB-231 cells transfected with individual siITGA2 #1-4 upon the scratched wound healing. B. Confirmation of reduced CD49b expression upon siITGA2 #1-4 mediated KD C. Images of slow filling of the scratched wound by MDA-MB-231 cells incubated with anti-ITGA2 mAb.

**Supplementary Fig S5.**
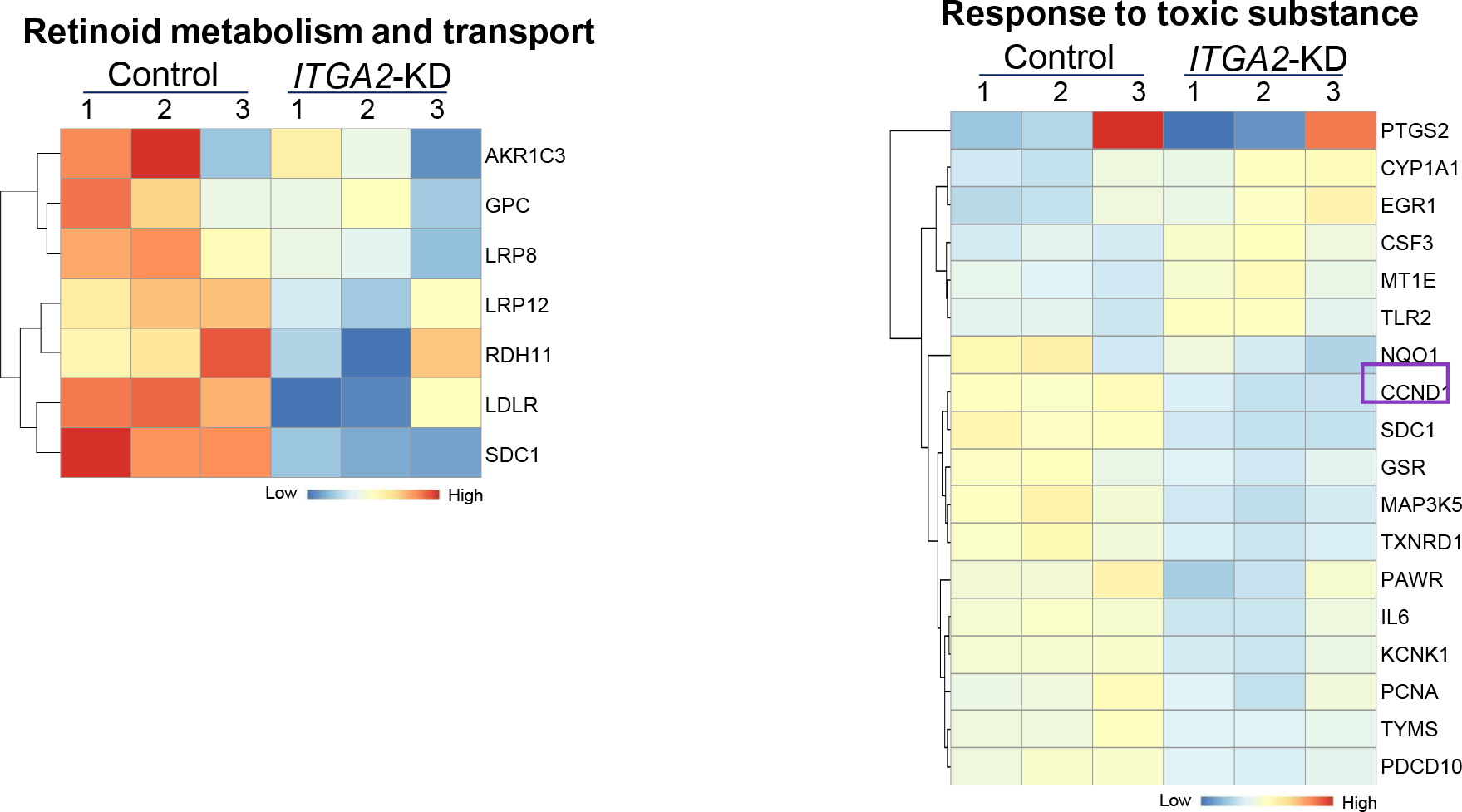
*ITGA2* KD-regulated pathways by RNA sequencing (related to Fig 5). Heatmap of genes in representative pathways of retinoid metabolism and transport, response to toxic substance, and metabolism of lipids.

**Supplementary Fig S6.**
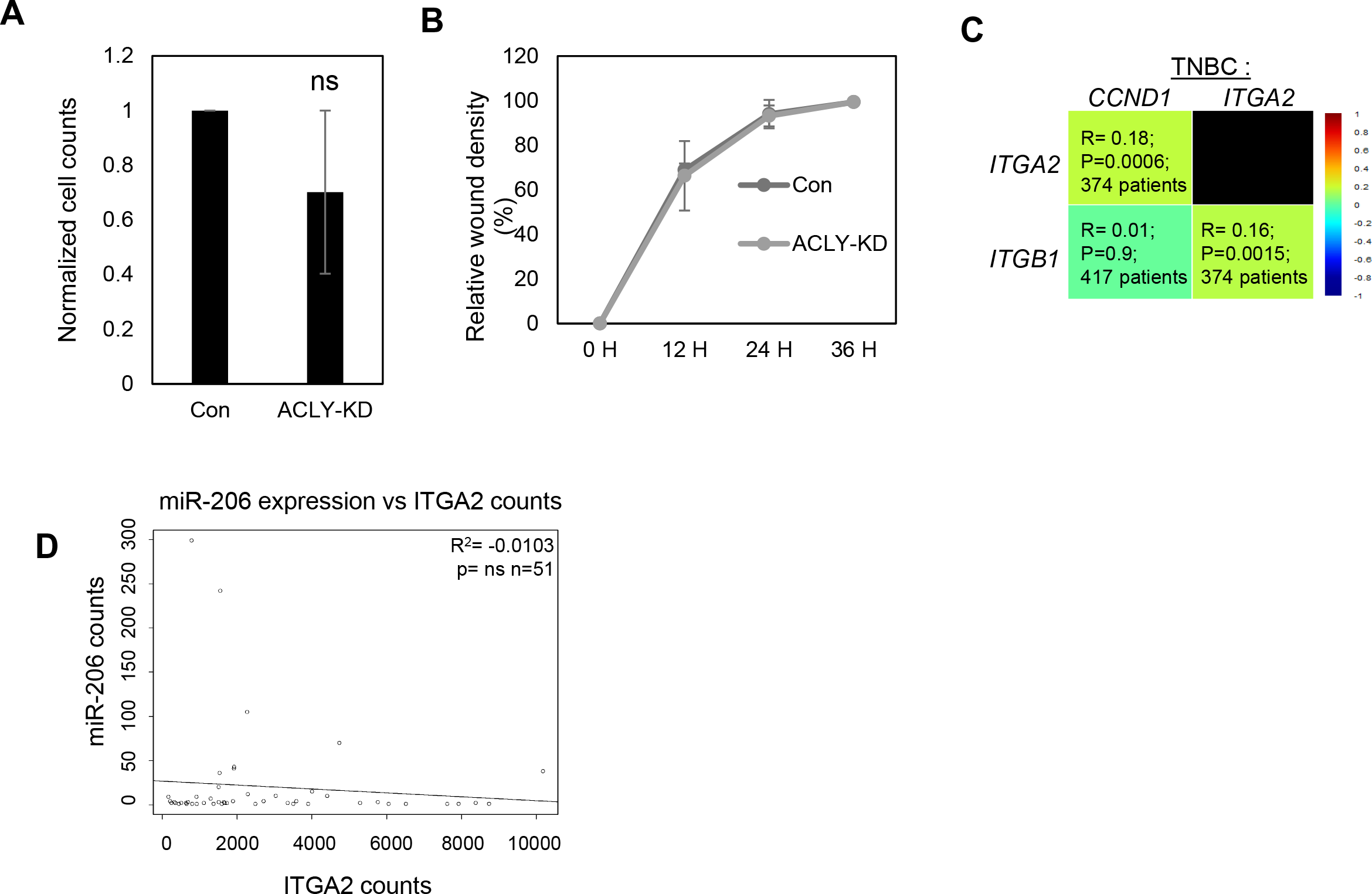
ACLY does not contribute to migration phenotype (related to Fig 6). A. No significant changes observed in *ACLY-*KD MDA-MB-231 cells. B. *ACLY* KD did not alter wound healing of MDA-MB-231 cells. C. Gene correlation analyses among *CCND1*, *ITGA2, and ITGB1* in TNBC using online Breast Cancer Gene Expression Miner v4.0. Significant associated P values and Pearson’s pairwise correlation coefficient R for the pairs of genes are shown in the boxes for *CCND1* and *ITGA2* as well as *ITGA2* and *ITGB1*. D. hsa-mir-206 expression and ITGA2 expression in triple negative breast cancer patients in TCGA database (n=51).

**Supplementary Fig S7.**
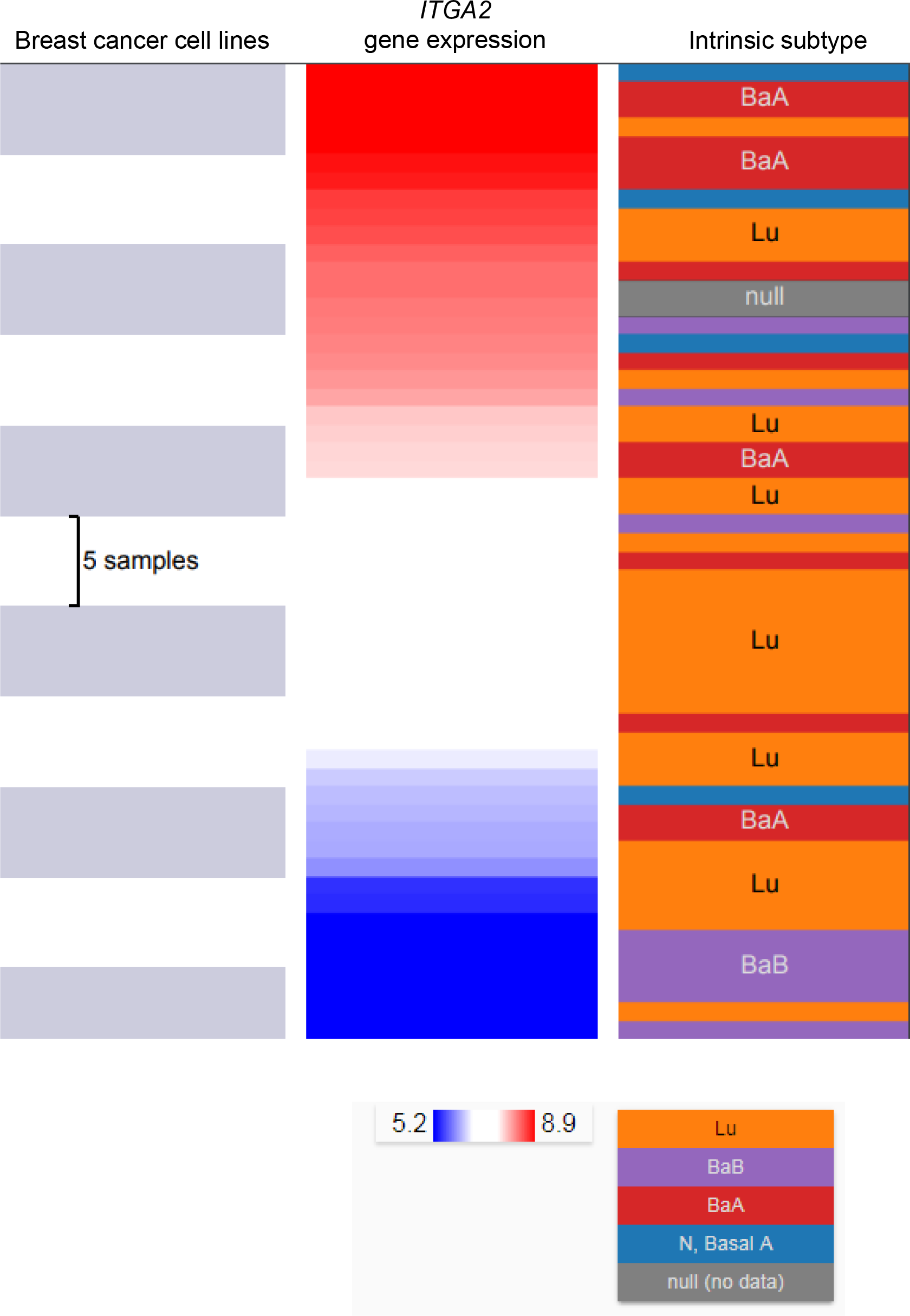
Expression of *ITGA2* in various breast cancer cell lines (related to Fig 7). UCSF Xena browser-based heatmap of ITGA2 expression across more than 50 cell lines and their corresponding origin of intrinsic subtype of breast cancer.

## References

1 Cancer, I. A. f. R. o. All cancers. The Glocal Cancer Observatory, online (2018).

2 Howlader N, N. A., Krapcho M, et al. SEER Cancer Statistics Review. (2016).

3 Siegel, R. L., Miller, K. D. & Jemal, A. Cancer statistics, 2018. CA Cancer J Clin 68, 7–30, doi:10.3322/caac.21442 (2018).

4 Anders, C. K. & Carey, L. A. Biology, metastatic patterns, and treatment of patients with triple-negative breast cancer. Clin Breast Cancer 9 Suppl 2, S73–81, doi:10.3816/CBC.2009.s.008 (2009).

5 Lapidot, T. et al. A cell initiating human acute myeloid leukaemia after transplantation into SCID mice. Nature 367, 645–648, doi:10.1038/367645a0 (1994).

6 Al-Hajj, M., Wicha, M. S., Benito-Hernandez, A., Morrison, S. J. & Clarke, M. F. Prospective identification of tumorigenic breast cancer cells. Proc Natl Acad Sci U S A 100, 3983–3988, doi:10.1073/pnas.05302911000530291100 [pii] (2003).

7 Diehn, M. et al. Association of reactive oxygen species levels and radioresistance in cancer stem cells. Nature 458, 780–783, doi:10.1038/nature07733 (2009).

8 Liu, H. et al. Cancer stem cells from human breast tumors are involved in spontaneous metastases in orthotopic mouse models. Proc Natl Acad Sci U S A 107, 18115–18120, doi:1006732107 [pii]10.1073/pnas.1006732107 (2010).

9 Zabala, M. et al. in Cancer Stem Cells (ed Justin D. Lathia) 25–58 (Academic Press, 2016).

10 Ramos, E. K., Hoffmann, A. D., Gerson, S. L. & Liu, H. New Opportunities and Challenges to Defeat Cancer Stem Cells. Trends Cancer 3, 780–796, doi:10.1016/j.trecan.2017.08.007 (2017).

11 Liu, X. et al. Homophilic CD44 Interactions Mediate Tumor Cell Aggregation and Polyclonal Metastasis in Patient-Derived Breast Cancer Models. Cancer Discov 9, 96–113, doi:10.1158/2159-8290.CD-18-0065 (2019).

12 Hsu, J. M. et al. STT3-dependent PD-L1 accumulation on cancer stem cells promotes immune evasion. Nat Commun 9, 1908, doi:10.1038/s41467-018-04313-6 (2018).

13 Kreso, A. & Dick, J. E. Evolution of the cancer stem cell model. Cell Stem Cell 14, 275–291, doi:10.1016/j.stem.2014.02.006 (2014).

14 Samaeekia, R. et al. miR-206 Inhibits Stemness and Metastasis of Breast Cancer by Targeting MKL1/IL11 Pathway. Clin Cancer Res 23, 1091–1103, doi:10.1158/1078-0432.CCR-16-0943 (2017).

15 Bockhorn, J. et al. MicroRNA-30c inhibits human breast tumour chemotherapy resistance by regulating TWF1 and IL-11. Nat Commun 4, 1393, doi:10.1038/ncomms2393 (2013).

16 Bockhorn, J. et al. Differentiation and loss of malignant character of spontaneous pulmonary metastases in patient-derived breast cancer models. Cancer Res 74, 7406–7417, doi:10.1158/0008-5472.CAN-14-1188 (2014).

17 Bockhorn, J. et al. MicroRNA-30c targets cytoskeleton genes involved in breast cancer cell invasion. Breast Cancer Res Treat 137, 373–382, doi:10.1007/s10549-012-2346-4 (2013).

18 Samaeekia, R. et al. MicroRNA-206 inhibits stemness and metastasis of breast cancer by targeting MKL1/IL11 pathway. Clin Cancer Res, doi:1078-0432.CCR-16-0943 [pii]10.1158/1078-0432.CCR-16-0943 (2016).

19 Liu, H. MicroRNAs in breast cancer initiation and progression. Cell Mol Life Sci 69, 3587–3599, doi:10.1007/s00018-012-1128-9 (2012).

20 Bartel, D. P. MicroRNAs: genomics, biogenesis, mechanism, and function. Cell 116, 281–297 (2004).

21 Ambros, V. The functions of animal microRNAs. Nature 431, 350–355, doi:10.1038/nature02871 (2004).

22 Desgrosellier, J. S. & Cheresh, D. A. Integrins in cancer: biological implications and therapeutic opportunities. Nat Rev Cancer 10, 9–22, doi:10.1038/nrc2748 (2010).

23 Adorno-Cruz, V. & Liu, H. Regulation and functions of integrin α2 in cell adhesion and disease. Genes and Diseases 6, In press (2019).

24 Gagliani, N. et al. Coexpression of CD49b and LAG-3 identifies human and mouse T regulatory type 1 cells. Nat Med 19, 739–746, doi:10.1038/nm.3179 (2013).

25 Mitra, S. K. & Schlaepfer, D. D. Integrin-regulated FAK-Src signaling in normal and cancer cells. Curr Opin Cell Biol 18, 516–523, doi:10.1016/j.ceb.2006.08.011 (2006).

26 Luo, M. et al. Mammary epithelial-specific ablation of the focal adhesion kinase suppresses mammary tumorigenesis by affecting mammary cancer stem/progenitor cells. Cancer research 69, 466–474, doi:10.1158/0008-5472.CAN-08-3078 (2009).

27 Yori, J. L. et al. Kruppel-like factor 4 inhibits tumorigenic progression and metastasis in a mouse model of breast cancer. Neoplasia 13, 601–610 (2011).

28 Carrer, A. et al. Acetyl-CoA Metabolism Supports Multistep Pancreatic Tumorigenesis. Cancer Discov 9, 416–435, doi:10.1158/2159-8290.CD-18-0567 (2019).

29 Stacey, D. W. Cyclin D1 serves as a cell cycle regulatory switch in actively proliferating cells. Current opinion in cell biology 15, 158–163 (2003).

30 Gkountela, S. et al. Circulating Tumor Cell Clustering Shapes DNA Methylation to Enable Metastasis Seeding. Cell 176, 98–112 e114, doi:10.1016/j.cell.2018.11.046 (2019).

31 Aceto, N. et al. Circulating tumor cell clusters are oligoclonal precursors of breast cancer metastasis. Cell 158, 1110–1122, doi:10.1016/j.cell.2014.07.013 (2014).

32 Szczerba, B. M. et al. Neutrophils escort circulating tumour cells to enable cell cycle progression. Nature 566, 553–557, doi:10.1038/s41586-019-0915-y (2019).

33 Guan, J. L. & Shalloway, D. Regulation of focal adhesion-associated protein tyrosine kinase by both cellular adhesion and oncogenic transformation. Nature 358, 690–692, doi:10.1038/358690a0 (1992).

34 Schlaepfer, D. D., Hanks, S. K., Hunter, T. & Geer, P. v. d. Integrin-mediated signal transduction linked to Ras pathway by GRB2 binding to focal adhesion kinase. Nature 372, 786, doi:10.1038/372786a0 (1994).

35 Ming, X.-Y. et al. Integrin α7 is a functional cancer stem cell surface marker in oesophageal squamous cell carcinoma. Nature Communications 7, 13568, doi:10.1038/ncomms13568 https://www.nature.com/articles/ncomms13568#supplementary-information (2016).

36 Bierie, B. et al. Integrin-beta4 identifies cancer stem cell-enriched populations of partially mesenchymal carcinoma cells. Proceedings of the National Academy of Sciences of the United States of America 114, E2337–E2346, doi:10.1073/pnas.1618298114 (2017).

37 Krebsbach, P. H. & Villa-Diaz, L. G. The Role of Integrin alpha6 (CD49f) in Stem Cells: More than a Conserved Biomarker. Stem cells and development 26, 1090–1099, doi:10.1089/scd.2016.0319 (2017).

38 Guan, J. L. Integrin signaling through FAK in the regulation of mammary stem cells and breast cancer. IUBMB Life 62, 268–276, doi:10.1002/iub.303 (2010).

39 Schwartz, M. A. & Assoian, R. K. Integrins and cell proliferation: regulation of cyclin-dependent kinases via cytoplasmic signaling pathways. Journal of cell science 114, 2553–2560 (2001).

40 Moreno-Layseca, P. & Streuli, C. H. Signalling pathways linking integrins with cell cycle progression. Matrix biology: journal of the International Society for Matrix Biology 34, 144–153, doi:10.1016/j.matbio.2013.10.011 (2014).

41 Roovers, K., Davey, G., Zhu, X., Bottazzi, M. E. & Assoian, R. K. Alpha5beta1 integrin controls cyclin D1 expression by sustaining mitogen-activated protein kinase activity in growth factor-treated cells. Molecular biology of the cell 10, 3197–3204, doi:10.1091/mbc.10.10.3197 (1999).

42 Wary, K. K., Mainiero, F., Isakoff, S. J., Marcantonio, E. E. & Giancotti, F. G. The adaptor protein Shc couples a class of integrins to the control of cell cycle progression. Cell 87, 733–743 (1996).

43 Ramirez, N. E. et al. The alpha(2)beta(1) integrin is a metastasis suppressor in mouse models and human cancer. The Journal of clinical investigation 121, 226–237, doi:10.1172/JCI42328 (2011).

44 Carrer, A. et al. Acetyl-CoA metabolism supports multi-step pancreatic tumorigenesis. Cancer Discov, doi:10.1158/2159-8290.CD-18-0567 (2019).

45 Dobin, A. et al. STAR: ultrafast universal RNA-seq aligner. Bioinformatics 29, 15–21, doi:10.1093/bioinformatics/bts635 (2013).

46 Anders, S., Pyl, P. T. & Huber, W. HTSeq--a Python framework to work with high-throughput sequencing data. Bioinformatics 31, 166–169, doi:10.1093/bioinformatics/btu638 (2015).

47 Love, M. I., Huber, W. & Anders, S. Moderated estimation of fold change and dispersion for RNA-seq data with DESeq2. Genome Biol 15, 550, doi:10.1186/s13059-014-0550-8 (2014).

48 Tripathi, S. et al. Meta- and Orthogonal Integration of Influenza “OMICs” Data Defines a Role for UBR4 in Virus Budding. Cell Host Microbe 18, 723–735, doi:10.1016/j.chom.2015.11.002 (2015).

49 Gyorffy, B. et al. An online survival analysis tool to rapidly assess the effect of 22,277 genes on breast cancer prognosis using microarray data of 1,809 patients. Breast cancer research and treatment 123, 725–731, doi:10.1007/s10549-009-0674-9 (2010).

50 Jezequel, P. et al. bc-GenExMiner: an easy-to-use online platform for gene prognostic analyses in breast cancer. Breast cancer research and treatment 131, 765–775, doi:10.1007/s10549-011-1457-7 (2012).

51 Jezequel, P. et al. bc-GenExMiner 3.0: new mining module computes breast cancer gene expression correlation analyses. Database: the journal of biological databases and curation 2013, bas060, doi:10.1093/database/bas060 (2013).

52 Mary Goldman, B. C., Akhil Kamath, Angela N Brooks, Jingchun Zhu, David Haussler. The UCSC Xena Platform for cancer genomics data visualization and interpretation. bioRxiv (2018).

53 Heiser, L. M. et al. Subtype and pathway specific responses to anticancer compounds in breast cancer. Proceedings of the National Academy of Sciences of the United States of America 109, 2724–2729, doi:10.1073/pnas.1018854108 (2012).

